# Hackathons as essential tools in an interdisciplinary biological training - report from trainings for Sub-saharan students

**DOI:** 10.1101/2025.03.27.645823

**Authors:** Marvin van Aalst, Tobias Pfennig, Faraimunashe Chirove, Marilyn Ronoh, Anna Matuszyńska

**Affiliations:** Computational Life Science, Department of Biology, RWTH Aachen University, Aachen, Germany; Cluster of Excellence on Plant Sciences (CEPLAS), Heinrich-Heine University Düsseldorf, Germany; Department of Mathematics and Applied Mathematics, University of Johannesburg, South Africa; Department of Mathematics and Statistics, University of Embu, Kenya

**Keywords:** authentic learning, project-based learning, datathon, modelling

## Abstract

Hackathons are collaborative, fast-paced events where participants from various fields work together to solve real-world problems. They are increasingly used in higher education to foster collaboration, problem-solving and applied computational skills, yet their role in interdisciplinary biological training remains under-documented. We report on the design and implementation of two computational biology summer schools in Kenya (2022, 2023), each culminating in a hackathon that integrated biological problems, quantitative methods, and coding. Both events targeted early-career researchers from multiple sub-Saharan countries and combined intensive teaching in programming and modelling with a time-bound group challenge using authentic marine conservation and synthetic epidemiological datasets. We describe the educational design, including its grounding in project-based learning, authentic learning, and Self-Determination Theory, and we document how performance-based assessment and structured participant feedback were used to evaluate learning outcomes. We present a critical reflective account of what worked, our teaching philosophy, and how hackathons can be embedded responsibly within biological curricula. We argue that, when embedded in sustained training and supported by appropriate mentoring, hackathons provide a practical and effective way to help biologists build computational skills, communicate across disciplines, and gain confidence in shaping their own research.

## 1 Introduction

### 1.1 Hackathons as a response to biology’s interdisciplinary data challenge

Modern biological research is constrained less by data availability than by the capacity to analyse and interpret complex datasets. High-throughput sequencing, imaging and environmental monitoring generate volumes of data [1]. To visualize the scale of this challenge, consider that 402.74 million terabytes of data are generated globally every day [2]; equivalent to handing out a smartphone, completely full of apps, pictures, messages, files, and video, to every third person on the planet, every day^1^. In biology specifically, technological advances in omics, imaging, and high-throughput screening have created an unprecedented data deluge. The Eli and Edythe L. Broad Institute of MIT and Harvard alone produces nearly 20 terabytes of genomic data daily [3].

However, making sense out of these data and answering relevant, disruptive questions that push the boundaries of biological knowledge requires more than computational power: it requires collaboration between experts in life sciences, mathematics, statistics and data science [4, 5]. Yet bringing these experts together is not simply about putting people in the same room and hoping they will identify both the problem and the solution. To effectively work together, we need to teach and cultivate a shared understanding and a common language, fostering smooth communication and collaboration at the intersection of diverse disciplines—an essential 21st-century skill emphasized by educational frameworks and workforce reports [6, 7]. This creates a pedagogical challenge for biology education: how do we train the next generation of biologists to work effectively across disciplinary boundaries? Traditional biology curricula, often structured around domain-specific knowledge delivered through lectures and laboratory practicals, may not adequately prepare students for collaborative, data-intensive problem-solving in authentic research contexts [8, 9].

### 1.2 Why hackathons for biological education?

Hackathons, intensive, collaborative events where participants work in teams to solve real-world problems within a constrained timeframe, offer a promising pedagogical approach for addressing this gap. The term combines “hack” (referring to creative, exploratory problem-solving, originating from the MIT Tech Model Railroad Club’s culture of clever technical solutions) and “marathon” (symbolizing the intensive, time-bound nature of the activity). While hackathons originated in software development, with the first recorded event organized by OpenBSD in 1999 [10], they have since expanded across disciplines [11–15], demonstrating particular value in contexts requiring rapid innovation and cross-disciplinary collaboration.

In biological education, hackathons hold specific promise because they mirror the authentic work of modern interdisciplinary research teams: they require participants to integrate diverse forms of expertise, communicate across disciplinary boundaries, work with real or realistic datasets, make decisions under time constraints, and produce tangible outputs [13]. Emerging evidence from engineering, software development, and health sciences demonstrates that hackathons can enhance problem-solving skills, teamwork, communication, and engagement [13, 16–18]. Recent systematic reviews confirm hackathons’ educational value across STEM disciplines [19, 20], though rigorous controlled studies remain limited [21], with few reports detailing how to design, implement, and assess hackathons as integral components of biology training programs [22].

### 1.3 Hackathons as capstone experiences in computational biology training

This report extends the evidence base for hackathons in biological education by presenting detailed accounts of two computational biology summer schools in Kenya (2022 and 2023), where hackathons served as capstone experiences for over 50 early-career researchers from five sub-Saharan countries. Both programs addressed a critical need: training biologists with little to no prior computational experience to work confidently with data and mathematical models, using authentic biological problems relevant to their regional contexts (marine conservation, HIV epidemiology, and harmful algal blooms).

Over the years, we, the authors, have first participated in, and later conducted, multiple hackathons in diverse settings. Our primary goal has been to enable students to use computational skills to address real-world biological problems. By fostering an environment that encourages open exchange of ideas and providing practical examples, we consistently observed a strong sense of achievement as students recognised their ability to tackle authentic challenges. While hackathons offer substantial opportunities for innovation and learning [23], their rapid pace and participant diversity can produce communication barriers, with differences in terminology, methods and disciplinary perspectives hindering collaboration. To mitigate these issues, we integrated hackathons not as isolated showcase events, but as the culminating components of structured training programs (summer school, workshop). Hackathons and workshops share a hands-on, active-learning foundation [24, 25], where hackathons emphasise rapid problem solving and knowledge integration [23], workshops offer structured, guided skill building. In combination, depth from workshops and the creative, solution-focused nature of hackathons reinforce each other, helping participants consolidate expertise while navigating interdisciplinary challenges in today’s data-driven biology.

This design allowed participants to consolidate newly acquired skills through authentic collaborative workflows while demonstrating competence in addressing real biological challenges. Hackathons are particularly well-suited to this capstone role because modern biology increasingly requires: (i) data literacy (importing, cleaning, visualizing, and interpreting complex datasets); (ii) computational thinking (translating biological questions into algorithmic or mathematical frameworks); (iii) interdisciplinary communication (bridging conceptual gaps between biology, mathematics, and computer science); (iv) applied problem-solving (working with incomplete information and real-world constraints); and (v) collaborative skills (integrating diverse expertise within teams). We present this work as a descriptive, analytically grounded experience report examining how pedagogical design choices shaped implementation and outcomes. Our goal is to contribute practical, theoretically informed guidance for biology educators seeking to incorporate authentic, collaborative problem-solving experiences into their programs, particularly in contexts where computational skills are underdeveloped and where bridging the gap between biological domain knowledge and data science capabilities is urgent.

## 2 Materials and Methods

### 2.1 Pedagogical Framework and Design Rationale

Our hackathon design was grounded in three complementary educational frameworks that shaped our implementation choices:

- **Project-Based Learning (PBL)**. We adopted PBL as our core instructional approach [26], where participants engage in hands-on discovery and problem-solving to create tangible outputs rather than passively receiving information [17]. Specifically, our hackathons required teams to: (i) formulate biologically meaningful questions from authentic and synthetic datasets, (ii) select and apply appropriate analytical methods independently, (iii) iterate their approaches based on feedback, and (iv) communicate findings to specific audiences. This structure positions learners as active knowledge constructors rather than consumers, which research shows enhances retention and transfer of skills [24, 27].
- **Authentic Learning**. We deliberately centered both training programs on socially relevant, real-world problems using genuine datasets from local contexts—sea turtle bycatch data collected by fishermen (Case Study 1) and epidemiological challenges affecting East African communities (Case Study 2). Authentic learning theory posits that learning is most effective when situated in contexts that mirror the complexity and social embeddedness of professional practice [28]. In our case, this meant participants worked with actual conservation data that would inform local decision-making, rather than sanitized textbook examples. The requirement to present findings in Kiswahili to fishermen (Case Study 1) further emphasized the authentic nature of the communication challenge facing interdisciplinary researchers.
- **Self-Determination Theory (SDT)**. Our design choices directly addressed the three psychological needs identified by SDT as essential for intrinsic motivation: **competence, autonomy**, and **relatedness** [29]. We fostered competence through progressive skill-building during the training weeks, followed by immediate application during the hackathon where participants could demonstrate mastery. The intensive mentoring support (approximately 1:10 trainer-to-participant ratio, see below) provided structure that allowed participants to succeed at challenging tasks without becoming overwhelmed. We supported autonomy by allowing teams to choose their own analytical approaches, decide how to divide responsibilities, and select visualization methods without prescriptive instructions. Relatedness was cultivated through deliberate team formation between disciplines and institutions, collaborative problem-solving structures, and social activities designed to build trust and community (poster sessions, field trips, informal discussions). This combination has been shown to enhance both learning outcomes and persistence in STEM fields [29–31].

These frameworks are interconnected: PBL provides the instructional structure, authentic learning ensures biological and social relevance, and SDT explains why this combination enhances motivation and engagement. Our “No One Left Behind” teaching philosophy (detailed below) operationalized these principles through specific design features.

### 2.2 Context, Participants, and Selection

#### Case Study 1: Summer School 2022, Watamu, Kenya

The first event was a two-week intensive program (September 5-16, 2022) titled “Computational Biology for Sustainable Environmental Research,” co-organized by RWTH Aachen University (Germany) and Mawazo Institute (Kenya). The program was held in Watamu, Kenya, a coastal town adjacent to one of the world’s oldest marine protected areas, providing immediate access to conservation sites and local environmental organizations.

##### Environmental Hackathon with real data

*Recruitment and selection*. We received 98 applications from 12 Sub-Saharan countries (Benin, Cameroon, Congo DRC, Kenya, Eswatini, Ethiopia, Ghana, Senegal, Somalia, South Africa, Tanzania, and Zimbabwe). The application required: (i) a 90-second video explaining motivation and research interests, (ii) a résumé, and (iii) written responses to three questions addressing current research, expected benefits from the training, and views on sustainable research. Five trainers independently evaluated applications using a rubric assessing research alignment, motivation, potential to benefit from and contribute to the program, and commitment to environmental sustainability, resulting in a selection of 25 participants.

*Participant characteristics*. The final cohort included 25 participants (72% women, 28% men; Fig. 1B) from six countries, with the majority (56%) residing in Kenya and significant representation from Ghana (32%), Uganda (8%), and Nigeria (4%). Educational levels ranged from undergraduate (8%) to post-doctoral researchers (4%), with the majority being Master’s students (64%) or doctoral candidates (20%) (Fig. 1C). Disciplines represented included marine biology, food security, environmental science, and policy. Critically, most participants reported little to no prior programming experience, though all possessed foundational knowledge in their biological domains. English proficiency was confirmed during video review; all instruction and materials were delivered in English, though participants presented hackathon findings in Kiswahili to local stakeholders.

*Funding and accessibility*. Participation was fully funded, covering accommodation, meals, all training materials, and travel. We provided travel grants to all participants who requested support, ensuring financial barriers did not limit access. Recognizing that 12% of applicants indicated childcare needs, we allocated budget for on-site childcare support; two participants used this service, bringing young children to the program.

**Fig 1.**
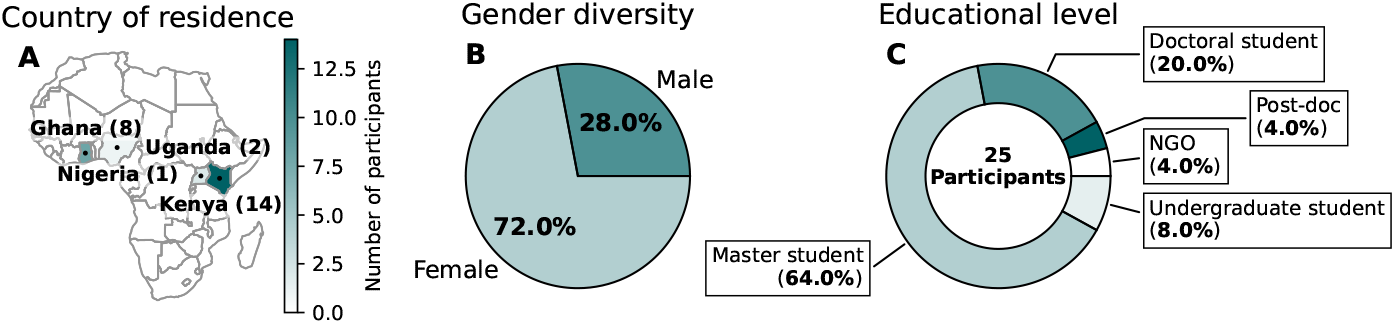
Participant distribution by country, gender, and education level in Case Study 1. A) Countries of residency of the 25 Sub-Saharan participants are marked on the map. The countries’ colors reflect the number of participants. B) The participants’ gender distribution. C) The educational level of the participants at the time of the hackathon. Participants stemmed from African universities or Non-Governmental organizations (NGOs).

#### Case Study 2: Workshop 2023, Embu, Kenya

The second event was a condensed five-day workshop (dates in September 2023) at the University of Embu, Kenya, focusing on computational modeling for epidemiology and environmental science.

##### Epidemiological Hackathon with generic data

*Recruitment and selection*. We distributed an open call through African university networks and social media, receiving applications from students across the continent. The selection process followed a similar rubric to Case Study 1, prioritizing applicants who would directly benefit from computational training for their ongoing research, resulting in a selection of 22 participants.

*Participant characteristics*. The cohort included 22 participants (63.6% women, 36.4% men; Fig. 2B) predominantly from Kenyan universities (91%), with one participant from Zimbabwe (4.5%) and Uganda (4.5%) (Fig. 2A). Educational levels were concentrated among Master’s students (72.7%) and doctoral candidates (22.7%), and undergraduate (4.5%) (Fig. 2C). Disciplines included mathematics, statistics, public health, and ecology. As in Case Study 1, most participants had limited programming experience but strong quantitative and biological foundations.

**Fig 2.**
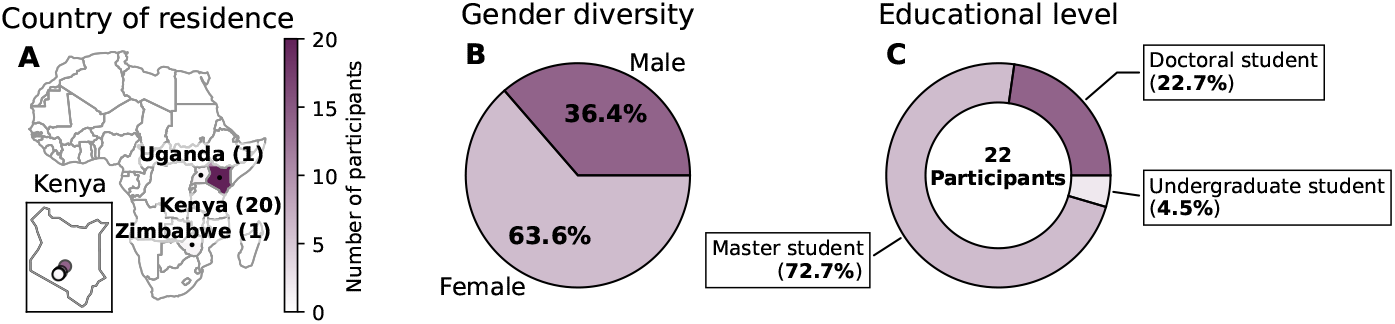
Participant distribution by country, gender, and education level in Case Study 2. A) Countries of residency of the 23 participants are marked on the map. The countries’ colors reflect the number of participants. The inset plot shows the location of the participants’ Kenyian universities. B) The participants’ gender distribution. C) The educational level of the participants at the time of the hackathon.

#### Limitations of sampling approach

We acknowledge several important limitations. First, both cohorts were self-selected from applicants motivated enough to complete a competitive application, potentially inflating apparent engagement and satisfaction relative to compulsory course settings. Second, the competitive selection process identified participants already primed for success—motivated, academically strong, and institutionally supported—limiting generalizability to broader student populations. Third, prior skill heterogeneity was substantial: while most had minimal coding experience, some had completed introductory programming courses or used computational tools in their research, creating variation in starting points that we accommodated through intensive mentoring but could not fully control. Fourth, language may have presented barriers for some participants, as technical instruction in English required fluency that, while verified, may have disadvantaged non-native speakers. Finally, our intensive support model (described below) is resource-intensive and may not be replicable in standard institutional settings. These factors mean our results reflect “best-case” conditions for hackathon implementation rather than typical classroom constraints.

### 2.3 Hackathon Structure and Implementation

Both hackathons followed a five-component structure (Fig. 3), though with different durations and data sources:

**Fig 3.**
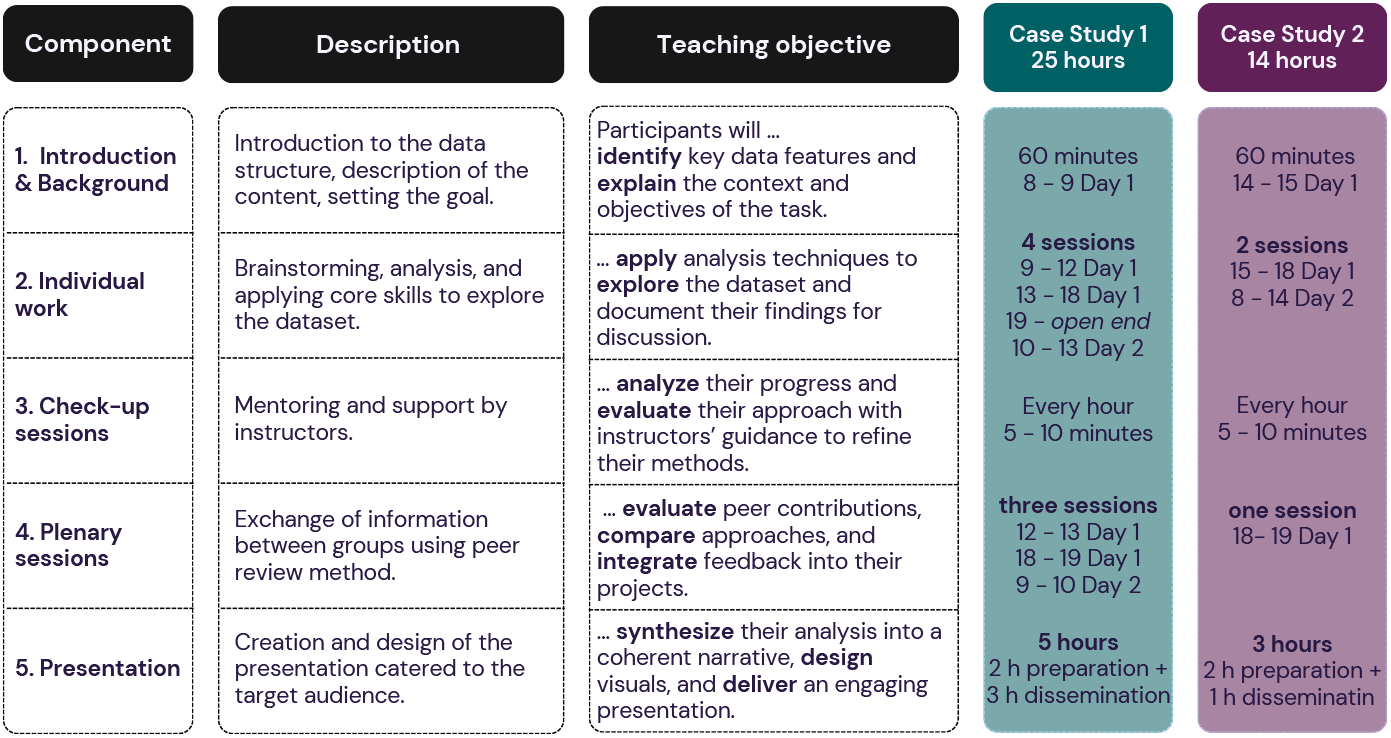
Structure of a Hackathon. Recommendation for organizing and holding a successful hackathon for education. Ensure that five critical components are present, regardless of the length of the planned event, starting from providing the context for the challenge and finishing with a proper dissemination of the final results to the most relevant stakeholders. The proposed teaching outcomes of each component are outlined according to Bloom’s Taxonomy of Educational Objectives for Knowledge-Based Goals [32] and the exemplary allocation of time is provided based on the experience from the two presented here case studies.

1. **Introduction and Context** (1 hour). Trainers introduced the dataset structure, biological/social context, and challenge objectives. In Case Study 1, Local Ocean Conservation (LOC) staff presented their turtle bycatch monitoring program and conservation goals. In Case Study 2, we framed the HIV and algal bloom challenges using recent literature and regional health/environmental statistics.
2. **Individual Group Work** (Case Study 1: 16 hours over 2 days; Case Study 2: 11 hours over 2 days). Participants worked in assigned teams (4 groups of 5-6 members each) on data exploration, analysis, and model development. Teams were deliberately heterogeneous, balancing: (i) disciplinary background (e.g., pairing marine biologists with statisticians), (ii) career stage (mixing undergraduates, Master’s, and PhD students), (iii) institutional affiliation, and (iv) geographic origin to maximize diverse perspectives and minimize pre-existing collaboration patterns. Teams had autonomy to choose their analytical approaches, division of labor, and presentation strategies.
3. **Mentoring Check-ins** (every 60 minutes, 5-10 minutes each). Trainers circulated continuously and conducted brief check-ins with each team every hour to: (i) assess progress and identify bottlenecks, (ii) provide technical troubleshooting (e.g., debugging code, clarifying statistical methods), (iii) offer conceptual guidance (e.g., suggesting alternative visualizations, prompting biological interpretation), and (iv) ensure equitable participation within teams. Mentors explicitly avoided prescribing solutions, instead using Socratic questioning to guide teams toward their own insights.
4. **Plenary Peer Exchange Sessions** (Case Study 1: 3 sessions; Case Study 2: 1 session). Teams briefly presented interim findings to the full cohort, received peer feedback, and coordinated to avoid duplicated efforts. This structure encouraged comparison of approaches and integration of complementary insights.
5. **Final Presentation Preparation and Dissemination** (Case Study 1: 5 hours; Case Study 2: 3 hours). Teams synthesized analyses into coherent narratives, designed visualizations, and prepared presentations tailored to their audiences. In Case Study 1, presentations were delivered in Kiswahili to local fishermen at Mida Creek, without electronic aids, requiring teams to create flipchart summaries accessible to non-technical audiences. In Case Study 2, teams presented to university leadership and peers using slides and live code demonstrations.

#### “No One Left Behind” Teaching Philosophy

Throughout both programs, we implemented an intensive support model to ensure accessibility for participants with limited computational backgrounds.

- **Extensive hands-on practice:** 75% of instructional time allocated to applied problem-solving rather than passive listening.
- **High trainer-to-participant ratio:** approximately 1 trainer per 10 participants, with at least 2 trainers present at all times during teaching blocks.
- **Frequent checkpoints:** trainers confirmed that all participants could reproduce examples and complete exercises independently on their own computers before proceeding.
- **Peer mentoring:** we encouraged faster learners to assist peers, fostering collaborative rather than competitive dynamics. **Technical troubleshooting support:** junior trainers (van Aalst, Pfennig) provided continuous one-on-one debugging assistance.

This model required substantial staffing (6 trainers for 25 participants in Case Study 1; 4 trainers for 22 participants in Case Study 2) but was essential for ensuring participants with no prior coding experience could fully engage.

### 2.4 Evaluation and Data Sources

To evaluate learning and the educational value of the hackathons, we used three complementary data sources aligned with the project-based design: (i) performance-based assessment of team outputs, (ii) structured participant feedback, and instructor observations. We did not use formal pre/post knowledge tests; instead, we treated the hackathon projects themselves as the primary demonstration of achieved outcomes.

First, each team’s project was assessed using a concise, pre-defined rubric applied to their learning artifacts/materials (Git repositories, notebooks, slides, outreach materials):

1. **Completion and coherence** – delivery of a functioning, self-contained project, including executable code or notebooks and a final presentation or summary.
2. **Use of taught methods** – appropriate application of core skills covered in the training (e.g. data import, cleaning, visualisation, basic modelling or statistical exploration) without major methodological errors.
3. **Biological relevance and interpretation** – formulation of a biologically meaningful question (e.g. turtle bycatch patterns, infection or bloom dynamics) and interpretation of outputs in a way that is consistent with the underlying biology and dataset limitations.
4. **Communication and audience alignment** – clarity and structure of the narrative for the intended audience (peers, local stakeholders, or university leadership), including where applicable the ability to translate technical results into accessible language.

We mapped these criteria to Bloom’s Taxonomy’s cognitive levels [32]:

- Criterion 1 (Completion) reflects *Application* (applying learned procedures)
- Criterion 2 (Methods use) spans *Application* and *Analysis* (selecting and implementing appropriate techniques)
- Criterion 3 (Biological interpretation) requires *Analysis* and *Evaluation* (interpreting data within biological context, judging validity)
- Criterion 4 (Communication) demands *Synthesis/Create* (reorganizing findings into coherent narratives for specific audiences).

The teaching team jointly reviewed all group outputs against these criteria to determine whether each project met the expected standard for successful completion. This performance-based evaluation served as our primary indicator of whether participants were able to integrate the computational tools and biological concepts addressed in the schools.

Second, immediately after each event, participants completed an anonymous online survey including Likert-type items on organisation, support, perceived learning gains, confidence in applying computational methods to biological problems, and open-ended questions on strengths and weaknesses of the format (see Fig. 4). The instrument was developed specifically for these events and is not a validated scale; we use these self-reports only as supportive, descriptive evidence and recognise their susceptibility to social desirability and novelty effects.

**Fig 4.**
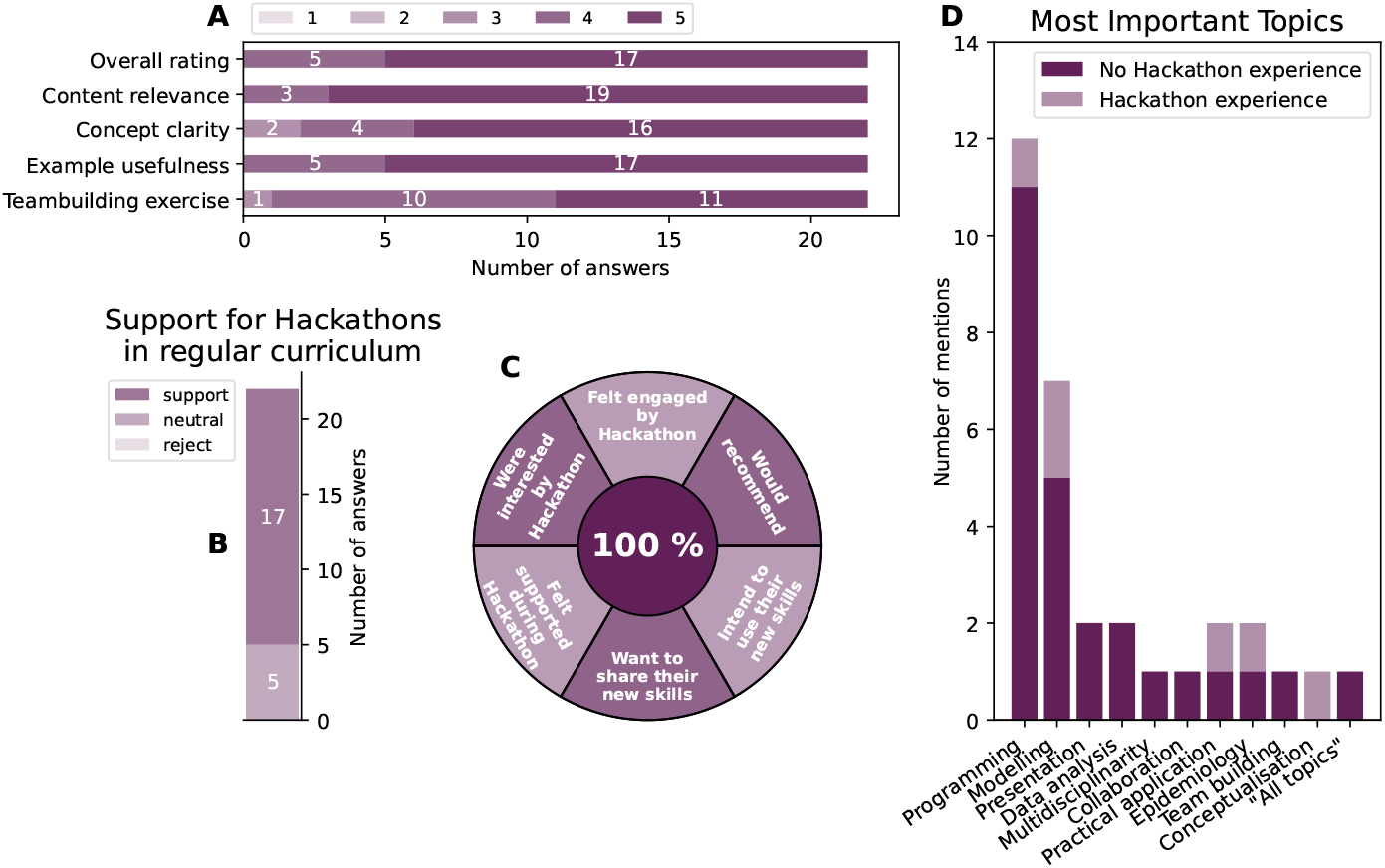
Participants’ evaluation results of Case Study 2. A) Participants’ rating of the hackathon-style workshop on a scale of 1 (worst) to 5 (best). B) Participants’ response to having Hackathons included in their regular curriculum. Answers with no clear position were counted as “neutral”. C) Summary of binary questions to the participants that were uniformly answered with “Yes.” D) The most participants’ important topics in the workshop, separated by whether they had previously participated in a hackathon. The participants’ free text answers were assigned all common topics explicitly mentioned. Figure D shows the number of mentions of each topic across participants. Results in B and D stem from free text (see Tables S2.

Third, instructors maintained brief field notes documenting observable patterns during the hackathons (engagement, collaboration, difficulties, reliance on mentoring, inclusion of less experienced participants). These notes were used to contextualise the performance and survey data and to identify recurring challenges and design implications. All analyses are descriptive; no inferential statistics are reported, and we do not claim causal proof of effectiveness.

## Data and materials availability

All teaching materials, participant projects, and documentation are publicly archived to support reproducibility and adoption by other educators:

Case Study 1 Summer School 2022: Teaching materials, schedules, and anonymized outputs are available at: https://git.rwth-aachen.de/computational-life-science/summer-school

Case Study 2 Workshop 2023: Complete teaching materials, Jupyter notebooks, participant project repositories, and workshop documentation are available at: https://github.com/Computational-Biology-Aachen/HackathonEmbu2023

These repositories include setup instructions (environment configuration, required software, installation guides), teaching notebooks with worked examples, and datasets used during training. We provide these resources to facilitate replication and adaptation by other institutions.

## 3 Results

### 3.1 Performance-based outcomes (primary evidence)

Using the pre-specified rubric (Methods), all teams in both case studies delivered a coherent, runnable project that:

1. **Completed a self-contained workflow** (executable notebooks/code and a final presentation).
2. **Applied taught methods** (data import/cleaning, visualisation, and, where relevant, basic modelling) without major methodological errors.
3. **Addressed a biologically meaningful question** and provided interpretations consistent with data limitations.
4. **Communicated findings** to the intended audience (stakeholders in Watamu; university peers/leadership in Embu).

In **Case Study 1**, teams identified spatial and gear-specific patterns in turtle bycatch and produced low-tech outreach artifacts for local fishers. In **Case Study 2**, teams formulated simple dynamical models (e.g., compartmental or growth models) for HIV-infected T cells and phytoplankton, related parameters to mechanisms, and contrasted dynamical behaviours across the two contexts. Taken together, artifacts indicate progression beyond *apply* towards *analyse/evaluate* and, in several cases, *create* levels on the revised Bloom taxonomy.

### 3.2 Participant perceptions (supporting evidence)

Post-event surveys (anonymous; non-validated, descriptive) indicated strong perceived value of the hackathon format. In **Case Study 2** (Fig. 4):

- Overall ratings for content relevance, concept clarity, and example usefulness were predominantly at the top of the scale.
- Binary items (engagement, recommendation, intention to apply/share skills) were uniformly positive.
- Free-text responses most frequently highlighted practical application, data analysis, programming, and teamwork as the most valuable topics; most comments supported including hackathons in the regular curriculum (Table S2).

We interpret these results cautiously (potential social desirability, novelty, and response biases) and use them to contextualise, not replace, the performance-based outcomes.

### 3.3 Process observations and inclusion

Instructor field notes documented high engagement across teams, frequent peer mentoring, and effective use of scheduled checkpoints to surface bottlenecks (e.g., debugging, choice of visualisations, framing biological questions). Heterogeneity in prior coding experience required sustained mentor presence (trainer:participant ∼1:4 to 1:10) to keep all participants progressing (“no one left behind”). This support intensity was integral to feasibility in both settings but is resource demanding. :contentReference[oaicite:3]index=3

### 3.4 Limitations (result-level)

Findings reflect best-case conditions: motivated, self-selected participants; short, intensive formats; and high mentor availability. There were no standardised pre/post tests or longitudinal follow-up; thus, we do not claim causal learning gains or skill retention. The evidence base for effectiveness in typical compulsory courses remains limited and warrants targeted evaluation in future iterations.

### 3.5 Summary

Across two implementations, all teams produced reproducible analyses addressing authentic biological questions and communicated results to appropriate audiences, meeting the primary performance criteria. Perception data and observations converged with artifact quality, suggesting that embedded, mentored hackathons function as feasible capstones for interdisciplinary biological training, while highlighting the need for lightweight, prospective assessment additions in future runs.

## 4 Discussion

The outcomes of both hackathons suggest that, when embedded within structured training programs and supported by close mentoring, hackathons can function as effective project-based capstones for interdisciplinary biological education. All teams delivered coherent, runnable projects that applied core methods to authentic biological questions, reaching at least the “analyse–create” levels of Bloom’s taxonomy. However, our evaluation approach (performance-based judgement of artifacts supplemented by self-reports and instructor observations) remains methodologically modest and is constrained by strong self-selection, heterogeneous prior skills, and intensive staff support. We frame our contribution as a descriptive, analytically grounded experience report that documents implementation under specific conditions (intensive, residential, internationally funded programs for highly motivated Sub-Saharan African early-career researchers). This positions our work transparently within the emerging evidence base while acknowledging what we cannot claim about generalisability or causal effectiveness.

We therefore recommend that future implementations retain the hackathon as an authentic, product-focused assessment, but strengthen their evidential basis through minimal, targeted additions: (i) publishing a short, transparent rubric for project evaluation (covering correctness, biological relevance, reproducibility and communication) and applying it systematically across teams; (ii) integrating a brief pre/post micro-task aligned with key learning goals (e.g. selecting appropriate visualisations, identifying data quality issues, formulating biologically meaningful questions) to capture shifts in applied reasoning without overburdening participants; and (iii) optionally collecting a short follow-up report on whether and how participants used the acquired skills in subsequent coursework or research. Such measures are feasible within the constraints of intensive schools, respect the principles of project-based and authentic learning, and would move hackathon reports beyond enthusiasm and satisfaction towards a more robust, comparable evidence base for their role in modern biology education.

### 4.1 Positioning Within the Hackathon Education Evidence Base

Our descriptive approach reflects the field’s current state. Systematic reviews show no standardized hackathon setup has been exhaustively investigated, with most studies relying on perceived rather than assessed learning [20]. Scoping reviews identify lack of long-term impact research as critical gaps [33, 34]. We share these constraints but advance the field through: (i) detailed implementation accounts reviews identify as lacking [21, 35], (ii) performance-based assessment addressing over-reliance on satisfaction surveys [36], and (iii) extension to biology education where evidence lags engineering contexts [19].

### 4.2 Recommendations for Educators

Based on our experience implementing hackathons in two computational biology training programs, we offer the following evidence-based guidance for educators considering similar approaches. These recommendations aim to maximize learning while acknowledging resource constraints that many institutions face.

#### Design and Structure

(1) Embed hackathons as capstone experiences following structured skill-building (1-2 weeks or semester-long courses), not as standalone events [17]. (2) Prioritize authentic, locally relevant biological problems through partnerships with conservation organizations, health agencies, or research institutes—authentic contexts enhance motivation and transfer [28]. (3) Design for heterogeneous skills using tiered challenges, frequent checkpoints, and peer mentoring to accommodate varied prior experience [20].

#### Support Infrastructure

(4) Maintain adequate mentoring capacity (approximately 1:10 trainer-to-participant ratio) through teaching assistants, peer facilitators, or reduced cohort sizes—immediate technical support prevents compounding frustration. (5) Balance intensity with well-being by scheduling breaks, normalizing error, and limiting duration (14-25 hours over 2 days worked well) to prevent burnout while maintaining productive challenge [23].

#### Implementation Details

(6) Compose teams intentionally across disciplines, career stages, and institutions to maximize diverse perspectives and prevent dominance by subgroups [37]. (7) Prepare technical contingencies including offline documentation, pre-installed software, spare hardware, and platform alternatives for inevitable technical failures. (8) Establish clear data use agreements addressing ownership, anonymization, and ethical responsibilities when using partner data.

#### Assessment and Adaptation

(9) Integrate assessment from the start using transparent rubrics, brief pre/post tasks, and structured reflection prompts rather than retroactive evaluation [38]. (10) Adapt formats to institutional contexts by considering mini-hackathons in existing courses, extendeding timelines with milestones, or switching to virtual platforms when residential intensive programs are infeasible [39].

### 4.3 Broader Implications for Biology Education

Biology’s computational transformation demands graduates who can translate between biological questions and quantitative methods, collaborate across disciplines, communicate to diverse stakeholders, and navigate real-world data complexity [9]. Traditional lecture-dominated, discipline-siloed curricula inadequately prepare students for this reality [40]. Hackathons offer one pedagogical approach for developing these interdisciplinary competencies, particularly when embedded as capstone experiences consolidating prior learning through authentic application. They complement rather than replace foundational coursework, laboratories, and research apprenticeships—occupying a valuable niche for synthesis, collaboration, and communication skills that resist traditional assessment [7]. Our Kenya case studies demonstrate feasibility in resource-constrained Global South settings, where computational training can build local capacity to address regional biological challenges (infectious diseases, biodiversity loss, climate impacts). The competence demonstrated by participants (many programming for the first time) challenges deficit narratives, illustrating that structured opportunity and quality instruction enable success [29]. However, pedagogical innovation requires rigorous evaluation. The field needs transparent reporting, performance-based assessment, longitudinal follow-up, comparative studies, and validated instruments enabling cross-study synthesis [20, 25, 41, 42]. As biology education evolves, we must couple experimentation with systematic evidence generation, building a robust knowledge base guiding future practice for data-intensive, interdisciplinary biological training.

## 5 Acknowledgments

This work was supported by the Volkswagen Foundation, UNESCO, the International Development Research Centre (Canada), and the Deutsche Forschungsgemeinschaft (DFG, EXC-2048/1 under Germany’s Excellence Strategy). We would like to thank the Local Ocean Conservation (LOC) in Watamu, Kenya, in particular Nicky Parazzi, who trusted us with their twenty-year-old data collection. The authors thank Julia Lorke and Tim Nies for critically reading the first draft of this article. Finally, we extend our gratitude to Otho Mantegazza, with whom we have created the educational content, for his involvement, ideas, and heart he put into educational programs, and to the Mawazo Institute, in particular Fiona Moejes, who was an essential co-organizer of the discussed above events.

## Disclaimer

In compliance with the Taylor & Francis AI Policy, we, the authors, disclose the use of ChatGPT (version 4) to enhance the clarity and coherence of our manuscript. This tool helped refine language and structure to effectively communicate complex ideas. We have thoroughly reviewed and take full responsibility for the content generated with ChatGPT’s assistance, ensuring its accuracy and integrity.

---

**Fig S1.**
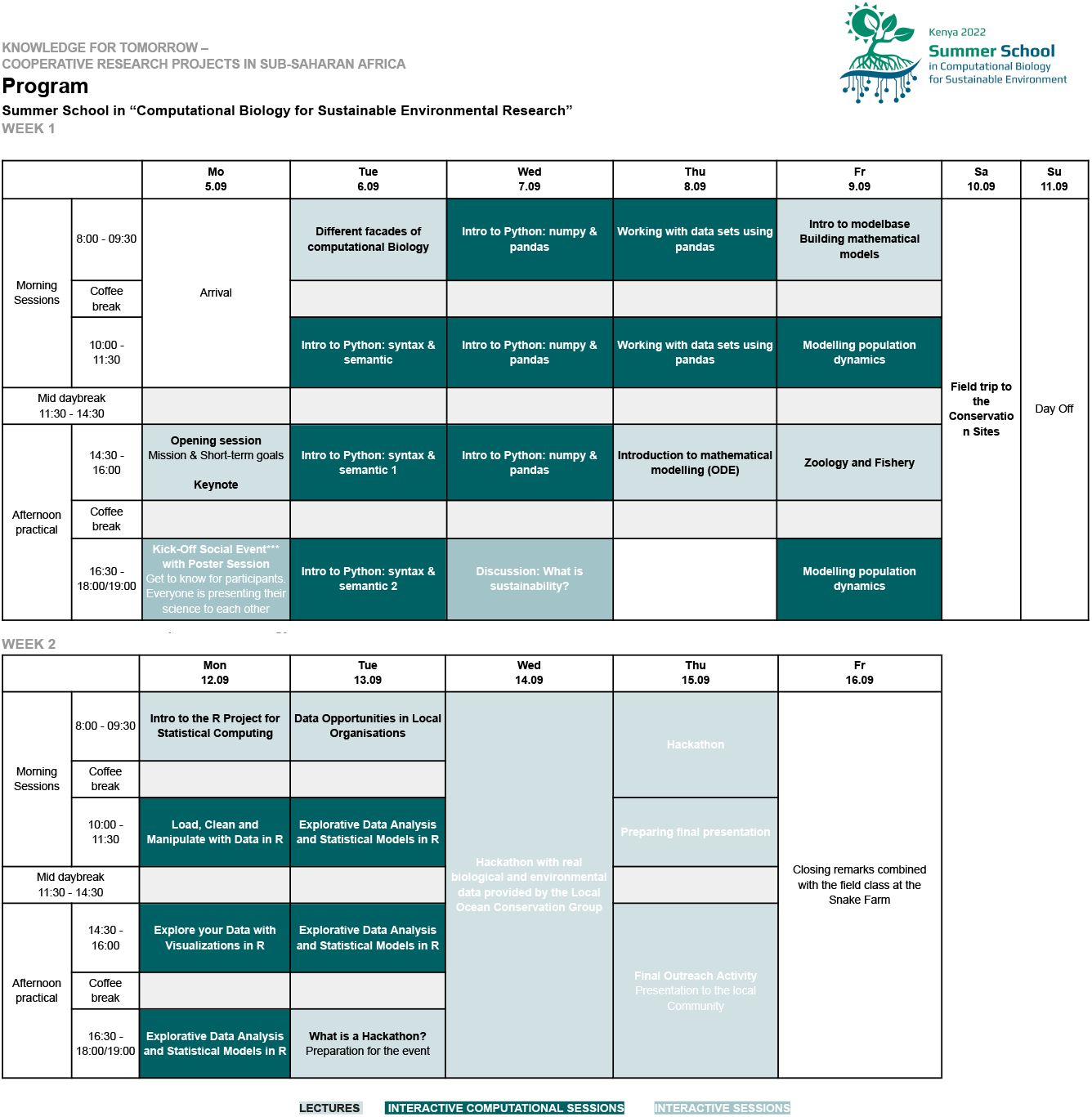
Case Study 1: Comprehensive Program Schedule for the Summer School in 2022, detailing the sequence of sessions, topics covered, and key activities designed to enhance participants’ learning experiences, including a 25-hours hackathon.

**Fig S2.**
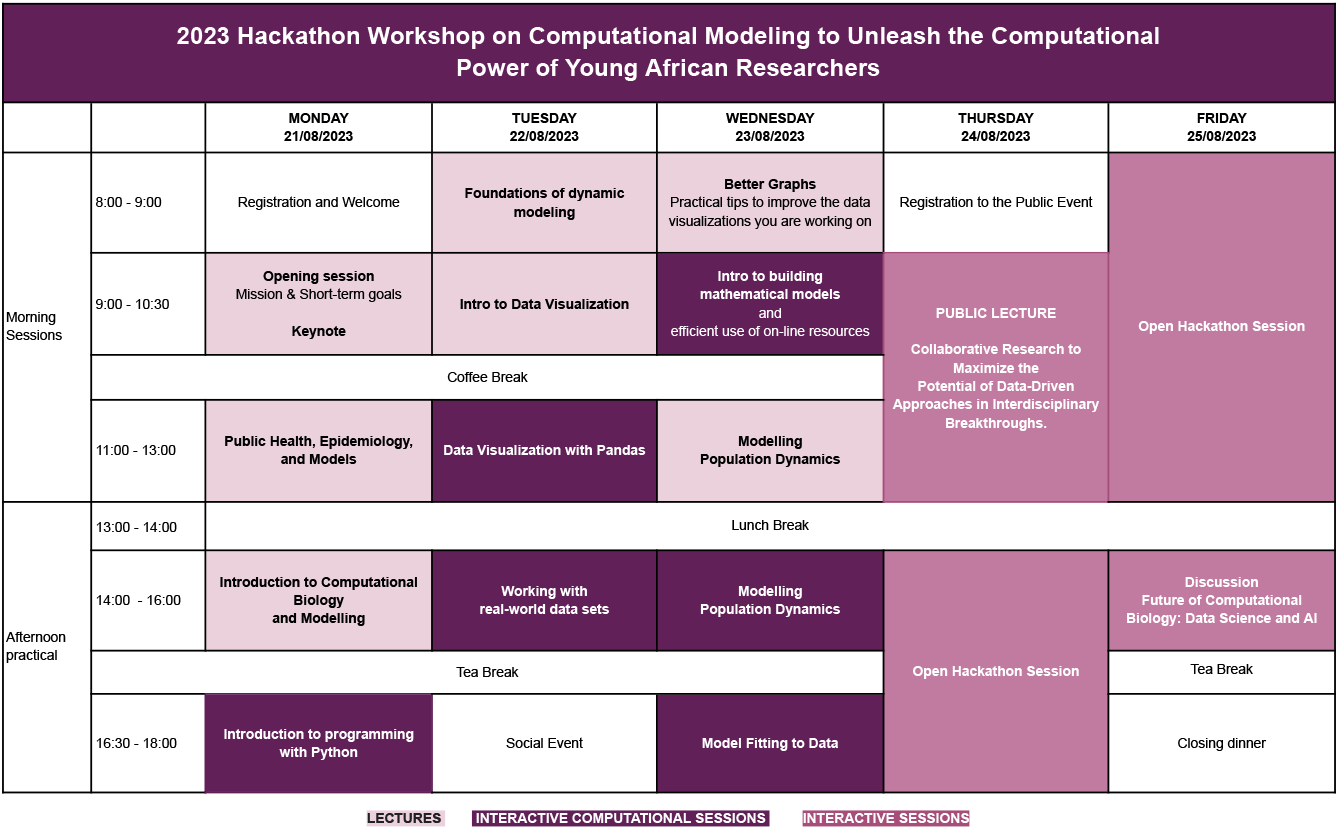
Case Study 2: Comprehensive Program Schedule for the Hackathon Workshop in 2023, detailing the sequence of sessions, topics covered, and key activities designed to enhance participants’ learning experiences, including a 14-hours hackathon.

**Table S2.**
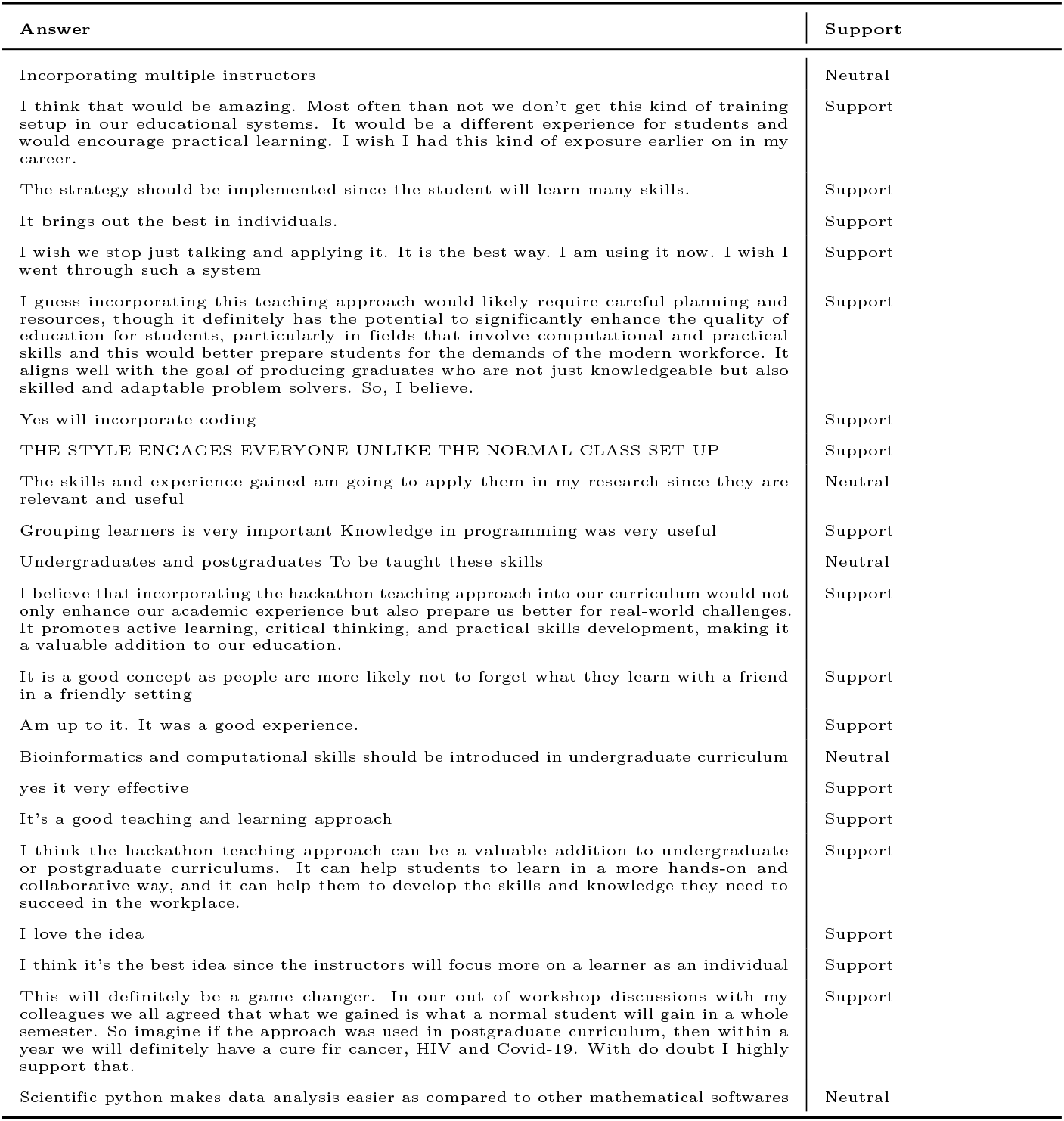
Summary of the open answer of the survey distributed among participants of the Case Study 2. Participants’ answers to the question “*After participating in this hackathon-style workshop, what are your opinions on incorporating the hackathon teaching approach into your undergraduate or postgraduate curriculum? This approach involves multiple instructors in a classroom setting, with hands-on practical activities following the introduction of each concept, including programming activities and group challenges to test the understanding of concepts and computational skills*.” The right column shows the interpreted support shown in Fig 4B. If an answer contained no explicit support for the incorporation of hackathons, it was counted as neutral.

1 assuming current population size to be 8.2 Billion and storage of a typical smartphone to be 128 GB

